# Seed banking alters native seed microbiome composition and function relative to natural populations

**DOI:** 10.1101/2024.07.16.603074

**Authors:** Dylan Russell, Vaheesan Rajabal, Matthew Alfonzetti, Marlien M. Van der Merwe, Rachael V. Gallagher, Sasha G. Tetu

## Abstract

- Seed banks are a vital resource for preserving plant species diversity globally. However, seedling establishment and survival rates from banked seeds can be poor. Despite a growing appreciation for the role of seed microbiota in supporting seed quality and plant health, our understanding of the effects of conventional seed banking processes on seed microbiomes remains limited.
- We investigated the composition and functional potential of the epiphytic seed microbiome of a native plant species using both 16S rRNA gene sequencing and culture-based approaches.
- Comparing the bacterial community composition of freshly collected seeds and those sourced from seed banking organisations, we found stored seeds hosted significantly less diverse bacterial populations, with substantial reductions in both low-abundance taxa and some core community members identified in unstored seeds. Bacteria with key plant growth promoting traits including IAA production, ACC deaminase activity, phosphate solubilisation, siderophore activity, and nitrogen fixation were identified in seed epiphytic communities, but these beneficial traits were less prevalent in stored seed compared to fresh seeds.
- Overall, these results suggest that epiphytic seed microbiomes may undergo significant changes during the storage process, selecting for bacteria tolerant to storage conditions, and potentially reducing the population of plant-growth promoting bacteria on seeds.

## INTRODUCTION

There has been a concerted global effort to preserve the seeds of plant species over the last three decades to safeguard against extinction and provide supply for in-situ conservation initiatives. It is standard for restoration seed suppliers and seed banking facilities, such as the Millenium Seedback (MSB) which holds ∼2.4 billion seeds, to reduce seed-moisture content to <10% and store in airtight containers at low temperatures (-18 °C to 15 °C) (Frischie et al. 2020). When appropriately stored, the viability of seeds from many species can be maintained for many years (DeVitis et al. 2020). However, seeds from such facilities may in some cases suffer poor rates of establishment in the field, hindering the effectiveness of this resource in meeting increasing demands (Merrit and Dixon 2011, Ceccon et al. 2016, Chapman et al. 2019). These inefficiencies have been attributed to a poor understanding of storage requirements, seed biology, and seed ecology (Streczynski et al. 2019, Turner et al. 2022), but it is not known whether alterations to seed-associated microorganisms during storage may also be important.

Microbial interactions with plants are known to significantly influence germination success and growth, forming the basis of an essential relationship in seed ecology (Vandenkoornhuyse et al. 2015, Nelson 2018). Seeds host a diverse array of fungi, bacteria, and archaea, including both endophytes and epiphytes, collectively forming the seed microbiome (Truyens et al. 2015, Simonin et al. 2022). These microbes can either be acquired from the surrounding environment or be maternally transmitted (Shade et al. 2017, Abdelfattah et al. 2023) and may include taxa with plant growth-promoting capacities (Berg and Raaijmakers 2018). Although the seed microbiome is typically less diverse than other mature plant organs (War et al. 2023), research indicates it serves as an advantageous reservoir of beneficial microbes that may contribute to the nascent seedling microbiome (Barret et al. 2015, Walsh et al. 2021, Arnault et al. 2024). Several studies demonstrate that seed-associated microorganisms can mediate seed dormancy and germination, fix and mobilise nutrients from the environment, regulate host hormones, and improve resistance to biotic and abiotic stressors (Goggin et al. 2015, Verma et al. 2018, Rodriguez et al. 2020, Hone et al. 2021, Matsumoto et al. 2021, Chimwamurombe et al. 2016). In light of this, the seed microbiome is anticipated to be a key determinant of host fitness during critical early stages through to maturity (Chesneau et al. 2020).

Contemporary understanding of seed microbiomes is mostly drawn from research in domesticated plant species, and knowledge of native seed microbiota remains limited (Mertin et al. 2023). As reviewed by Simonin and colleagues (2022), seeds are known to host a core microbiome shared among individuals and populations of a given host species (Shade et al. 2017) which is suggested to be evidence of coevolved microbial symbioses fulfilling important biological and ecological functions in hosts (Wasserman et al. 2022). Commonly reported core genera such as *Pantoea*, *Pseudomonas*, *Sphingomonas*, and *Methylobacterium* are found in many plants, including native species (Simonin et al. 2022). Evidence suggests both endophytes and epiphytes may constitute the core seed microbiome (Links et al. 2014, Morales-Moreia et al. 2021, Nelson 2018), however the epiphyte community is more likely to be influenced by local environmental factors and reflect the surrounding microbial diversity (Rochefort et al. 2019, Klaedtke et al. 2016). It has also been recognised that native seeds tend to host more diverse microbial communities compared to domesticated species, and host taxa with well-documented plant-growth-promoting capacities (Wasserman et al. 2019, Kim et al. 2020, Chandel et al. 2022, Mertin et al. 2023).

Despite increasing recognition of the importance of the seed microbiome, the effect of storage practices remains largely unknown (Berg and Raijmakers 2018), particularly for non-agricultural species. Research to date has indicated that simulated storage conditions can result in the loss of bacterial diversity in seeds but these findings are primarily derived from agricultural host species using surface sterilised seeds, which removes the epiphytic fraction prior to analysis. Previous work by Chandel and colleagues (2021) on soybean [*Glycine max* (L.) Merr.] endophytes indicate the most significant reduction in diversity occurs during the initial drying stages, with further gradual losses occurring over time in storage. Additionally, temperature was identified to influence the rate of bacterial diversity loss during storage in both soybean and rice [*Oryza sativa* (L.)] seed endophytes (Chandel et al. 2021, Dutta et al. 2022). Overall, seed storage procedures appear to favour the retention of taxa tolerant to desiccation and low temperatures such as *Pantoea* and *Bacillus*, although this has not been reported for seed retrieved from seed-banking facilities.

To assess the impact of seed storage on the microbiome of species commonly used in ecological restoration, we compared the bacterial epiphytic seed microbiome from natural and stored populations of *Acacia ulicifolia* (Salisb.) Court. across the Greater Sydney Basin (New South Wales, Australia). Specifically, we compared the seed-associated microbiome from undisturbed natural populations to seven seed samples of *A. ulicifolia* obtained from seed banking organisations. Analyses considered both bacterial community composition and load, using a combination 16S rRNA gene community sequencing, flow cytometry, and culture-based investigations. Cultured isolates were also subject to laboratory assays to assess trait profiles associated with plant growth-promoting functions. Given the exposure of seed surfaces to the environment, we hypothesised native seed epiphytic communities to vary between populations, and between stored and freshly collected seeds. Additionally, we anticipated epiphytes associated with stored seed to possess adaptive strategies toward low temperature and low humidity environments.

## MATERIALS AND METHODS

### Seed collection and processing

Seeds for this study originated from two sources; freshly collected seed pods and seeds obtained from two seed banking organisations. Populations of *A. ulicifolia* were sourced by examining occurrence records for the species in the Atlas of Living Australia (https://www.ala.org.au/) (accessed September 2022). Freshly collected (hereafter fresh) seed pods were sampled from five separate locations (Table S1). From each location, three biological replicates of *A. ulicifolia* seed pods were collected while wearing gloves to avoid contamination. Each replicate consists of opened pods bearing seeds, collected from 3 to 5 individuals growing within a 10-meter radius and replicates were spaced at least 100 meters apart. All samples were collected under a Scientific Licence granted by the NSW Government. Stored *A. ulicifolia* seeds included in this study were sourced from two organisations routinely involved in seed banking and storage; Greening Australia (GA), and the Australian PlantBank (Botanic Gardens of Sydney, Mount Annan, NSW) (PB). Four accessions stored at 15 °C were obtained from GA, and three accessions stored at 4 °C were obtained from PB. Both groups of samples were stored according to standard methods, in airtight aluminium bags. All accessions were treated as independent samples in all subsequent experiments. Locations of seed sampling sites are indicated in Figure 1.

**Figure 1:**
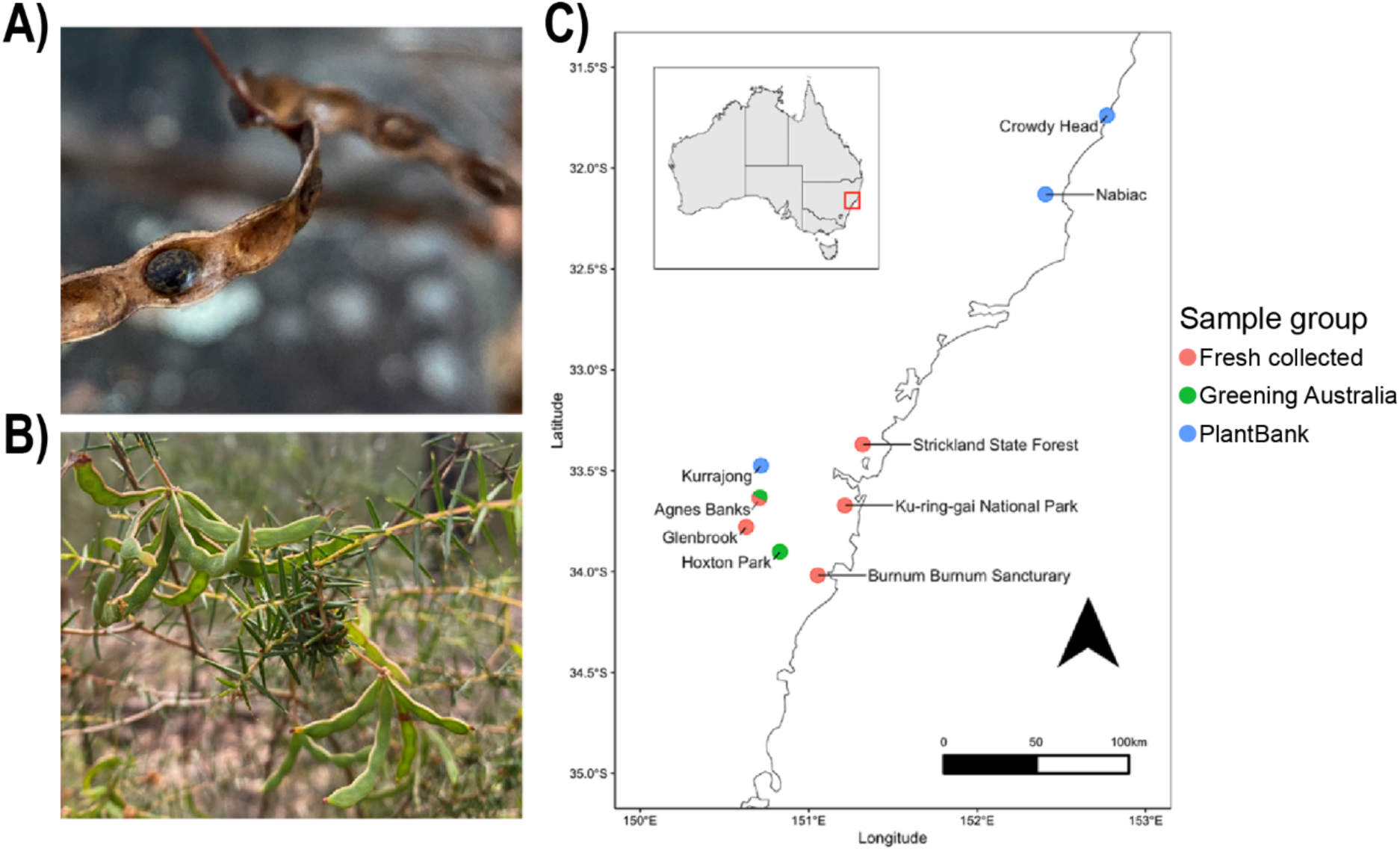
**(A)** Opened and **(B)** unripe unopened *A. ulicifolia* fruit. **(C)** Map of locations from which seed samples were obtained in this study. Sample collection group indicated by colour. *Orange*: Samples freshly collected from natural populations for this study. *Green*: sources of samples collected and later stored by Greening Australia. *Green/orange*: location where fresh collection and Greening Australia samples have shared geographic source. *Blue*: sources of samples collected and later stored by Australian PlantBank.

Seeds were extracted from freshly collected seed pods using flame-sterilised forceps. Per sample, 0.2 g (±0.1 g) of seed was added to 1 mL of sterile phosphate buffered saline with 0.05% Tween-20 (PBST) in a sterile microfuge tube. Epiphytes were collected by vortexing at 2500 RPM for 2 min, followed by sonication at 40 kHz for 30 min. Plant debris was removed by filtering supernatants through a 100 µm cell strainer (Corning, USA). Aliquots were taken and stored at -20 °C for DNA extraction, in addition, glycerol stocks of the epiphyte washes were prepared and stored at -80 °C, and the remainder of each sample was stored at 4 °C for culturing and flow cytometric analysis. Bacterial isolation and flow cytometric measurements from all epiphyte washes were performed within 24 hours of preparation to ensure results were as representative of populations *in planta* as possible.

### Bacterial isolation

#### Isolation of seed-associated bacteria

Epiphyte washes were serially diluted in sterile phosphate buffered saline (PBS) and plated onto nutrient agar (NA) plates amended with 2.5 µg mL^-1^ amphotericin B to suppress fungal growth. Plates were incubated at 28 °C for 4 days followed by 22 °C for 6 days. Colonies were counted, and those determined to be morphologically distinct were picked and subcultured onto fresh NA plates. Pure isolate cultures were grown in nutrient broth (NB), qualitatively screened for a range of putative plant growth-promoting phenotypes, and cryopreserved in 25% glycerol at -80 °C. Prior to functional characterisation, isolates were grown in NB and washed in sterile PBS by centrifugation 3 times to remove residual media. All glassware used in media preparation was cleaned with 6 M HCl and rinsed thoroughly with ddH2O prior to use.

#### Screening for selected plant growth-promoting phenotypes

##### Phosphate solubilisation capacity

Inorganic phosphate mobilisation was assessed by culturing isolates on National Botanical Research Institute’s phosphate growth medium (NBRIP) (Nautiyal 1999). First, 2 µL of isolate culture was spotted onto NBRIP agar in triplicate and incubated at 22 °C. Phosphate solubilisation capacity was determined by the development of a clear halo surrounding colonies after 7 days.

##### Nitrogen fixation

Biological nitrogen fixation was evaluated based on growth in nitrogen-free media. Prior to media preparation, all glassware was cleaned with 6 M HCl and rinsed thoroughly with ddH2O. Liquid NFb media was prepared as described by Cordova-Rodriguez et al. (2022). A 96-well plate containing 295 µL NFb was inoculated with 5 µL of isolate culture and incubated at 28 °C for 7 days. A colour change from green to blue was inferred to indicate nitrogen fixing capacity.

##### Biosynthesis of auxin (indole-3-acetic acid, IAA)

Auxin biosynthesis was assessed by the colorimetric method described by Khalaf and Riazada (2016). First, IAA production was induced by inoculating 290 µL of Luria-Bretani (LB) broth (Difco Laboratories, USA) supplemented with 5 mM L-tryptophan and incubated at 28 °C for 4 days. Cell-free supernatants were collected by centrifugation at 4800 x g for 15 minutes and transferred to a 96-well plate in triplicate. Equal volumes of Salkowski’s reagent (0.5 M FeCl3 in 35% HClO4) were added to each well and the reaction was incubated in the dark at 22 °C for 30 minutes. IAA production was measured by spectrophotometric measurement at 530 nm, relative to *Escherichia coli* DH5α and uninoculated culture controls.

##### Siderophore production

Siderophore production was estimated by liquid chrome azurol S (CAS) assay (Gu et al. 2021, Arora and Verma 2017). Briefly, iron-limited modified King’s B medium (MKB) was prepared in acid washed glassware as outlined by Gu et al. (2021). Next, the assay solution was prepared by dissolving 121 mg CAS in 100 mL ddH2O and 20 mL of 1 mM FeCl3 solution prepared in 10 mM HCl. This was slowly added to a 20 mL solution of 5 mM hexadecyl trimethyl ammonium bromide (HDTMA) while stirring. Isolates were grown in MKB media for 3 days at 28 °C, and cell-free supernatants were collected by centrifugation. Equal volumes of supernatant and CAS-HDTMA solution were transferred to a 96-well plate and following incubation for 30 min at room temperature, absorbance at 630 nm was measured using a spectrophotometer. Siderophore activity was determined by measuring a decrease in A630 relative to uninoculated controls.

##### ACC deaminase activity

Isolates were screened for 1-aminocylopropane (ACC) deaminase activity by the ninhydrin method (Li et al. 2011). First, Dworkin and Foster salts (DF) medium was prepared following the protocol of Penrose and Glick (2003). Next, solutions of DF media were prepared containing either 3 mM ACC or (NH4)2SO4 and inoculated with isolates in a 96-well plate. Following incubation at 28 °C for 4 days, culture supernatants were collected by centrifugation, and diluted 10-fold with fresh DF media. A ninhydrin reagent was prepared by dissolving 500 mg of ninhydrin and 15 mg of ascorbic acid in 60 mL of ethylene glycol and mixed with 60 mL of 1 M citric acid (pH 6.0) immediately prior to analysis. ACC deaminase activity was measured by adding equal volumes of dilute DF-ACC supernatant to the ninhydrin reagent followed by boiling for 30 minutes and measurement of absorbance at 570 nm, relative to uninoculated DF-ACC controls.

#### Taxonomic identification of isolates

Isolates with plant growth-promoting traits of interest were selected for taxonomic identification via polymerase chain reaction (PCR) and 16S rRNA gene sequencing using the universal primer set 27F (5’-AGAGTTTGATCMTGGCTCAG-3’) and 1492R (5’-TACGGYTACCTTGTTACGACTT-3’) which amplify close to the full length of the 16S rRNA gene (Frank et al. 2008). Amplicons were generated in 25 µL reactions using GoTaq 2X master mix (Promega, USA), 0.5 µM of each primer, and 0.75 µL template DNA. PCR was performed at 98 °C for 2 min, followed by 30 cycles of 98 °C for 30 sec, 55 °C for 30 sec, and 72 °C for 1.5 min, and a final extension at 72 °C for 10 min. Amplicons were assessed for size and purity by electrophoresis on a 2.0 % agarose gel, and once confirmed, purification and Sanger sequencing using 27F primers was carried out by Macrogen (Seoul, RoK). Sequences were trimmed based on basecalling quality and searched against the SILVA (v. 138) 16S rRNA gene database website using the Alignment, Classification and Tree (ACT) tool with a cut-off threshold of 0.97. Sequences were aligned and a phylogenetic tree was built based on maximum likelihood (RAxML).

### Flow cytometric analysis of epiphytic cell densities

The microbial cell densities of seed epiphyte washes were estimated by flow cytometry following well-established methods with minor modifications (Marie et al 1997, Focardi et al 2022). Samples were stained by incubating a 50 µL aliquot of epiphyte wash with a 1:10,000 dilution of SYBR Green I (SYBR) (Invitrogen) at 22 °C for 1 hour in the dark, PBS was passed through a 0.22 µm filter (Sartorius, Germany) and likewise stained to assess background noise. Stained samples were serially diluted 1:10 and 1:100 in 0.22 µm filtered PBS, and four technical replicates were prepared for each dilution. Sample analysis was performed using a Beckman CytoFLEX S flow cytometer (Beckman Coulter, USA) using a 488 nm laser for the analysis of forward scatter (FSC), side scatter (SSC), and SYBR fluorescence signal (525/40). The instrument was run using ddH2O as sheath fluid. Instrument calibration was performed prior to analysis using CytoFLEX Daily QC Fluorospheres (Beckman Coulter, U.S.A.). FSC trigger signal and thresholding were empirically determined by analysing a tube containing a mixture of Fluoresbrite yellow-green (YG) 1.00 µm polystyrene beads (Polysciences, USA) and CaliBRITE FITC 6 µm polymethacrylate beads (BD, USA). Based on the FSC signal intensity for YG beads (1050), the FSC trigger was set to 800 arbitrary units for all further analysis. Samples were acquired at a 10 µL min^-1^ flow rate for 2 min and gating was performed using the CytExpert (v 2.5) software. Gates were drawn to isolate singlet events within the 1-5 µm size range with SYBR fluorescence intensities greater than 100 au at 525 nm. Counts were imported to R (v 4.2.3) (R Core Team 2023) for background correction and statistical analysis.

## Bacterial community profiling

### DNA sequencing and library construction

DNA was extracted from all seed epiphyte washes using the DNeasy PowerSoil DNA extraction kit (Qiagen, Germany) following the manufacturer’s specifications. DNA quality and yield were assessed on a NanoDrop ND 1000 and a Qubit 3 fluorometer, respectively.

Amplicons of the near full-length 16S rRNA gene were generated by PCR using universal primers 27F and 1492R with unique custom barcodes. Reactions consisted of 25 ng template DNA, 10 µL 5x GC reaction buffer, 1 µL of 10 mM dNTP mix, 0.5 µL of each 50 µM primer, 1.5 µL DMSO, 0.25 µL Phusion polymerase (New England Biolabs, USA), and 30 µL NF-ddH2O (Invitrogen). PCR was performed at 98 °C for 2 min, followed by 35 cycles at 98 °C for 30 sec, 60 °C for 30 sec, and 72 °C for 1.5 min, and a final extension at 72 °C for 10 min. Reactions were prepared in triplicate and pooled. All samples were successfully amplified except for Kurrajong 1988; despite optimisation efforts, no product could be obtained for this sample due to low DNA yield, and so was not carried onto sequencing.

Barcoded amplicons were purified using AMPure XP beads (Beckman Coulter, USA), resuspended in 25 µL NF-ddH2O, and quantified using a Qubit fluorometer. An equimolar multiplexed 16S rRNA gene amplicon library was then generated using custom barcoded primers (Table S2) and the Oxford Nanopore Technologies (ONT) Ligation Sequencing Kit (SQK-LSK110) as per the manufacturer’s specifications. Sequencing was performed using a R9.4.1 flow cell and MinION MK1B device. Basecalling was performed using Guppy (v 6.3.2) using the high-accuracy basecalling model. Raw reads were exported for each barcode in FASTQ format.

### Bioinformatic analysis of community sequencing data

Demultiplexed raw reads for 16S rRNA amplicons were processed using the Spaghetti pipeline (Latorre-Perez et al. 2021). This process involved adapter removal by porechop, read filtering to retain reads with a size range of 1.2–1.8 kb using Nanofilt, quality control with Nanostat, chimera detection and removal using ycard, and read mapping against the SILVA (v. 138) database by mimimap2, to the species level. One sample from the STR sample group (STR1) was removed from the dataset due to the low number of reads recovered. Processed sequencing data were imported to R (v. 4. 2. 3) as taxonomy and abundance tables, and associated metadata. Host-associated mitochondria and chloroplast reads, and low abundance features (<10 counts) were removed from the dataset, and samples were rarefied to the lowest number of read counts (31,140 reads) without replacement using the phyloseq R package (v 1.44.0) (McMurdie and Holmes 2013). Next, operational taxonomic units (OTUs) were agglomerated to the species level to remove artificial heterogeneity which can be introduced into Nanopore sequencing data, as per Latorre-Perez et al. (2021). Data was exported to the microeco (v. 1.2.2) (Liu et al. 2021) package for all subsequent analyses using the file2meco package (v 0.5.0) (Liu et al. 2021).

Taxonomy-based functional prediction was performed using the Tax4Fun2 (Wemheuer et al. 2020) function from the microeco package and BLAST command line package (v 2.13.0) (Camacho et al. 2009). KEGG ortholog abundances obtained from Tax4Fun2 prediction were annotated using the PLaBAse plant growth promoting trait ontology (PGPTO) (Patz et al. 2021).

### Statistical analysis

All statistical tests and data visualisations were performed in R using the microeco and ggplot2 packages (Wickham 2016). Differences in Shannon diversity, Chao1 richness, and Inverse-Simpson (InvSimpson) evenness were tested for significance by the Kruskal-Wallis pairwise test. Beta diversity based on a Bray-Curtis dissimilarity matrix was assessed by principal coordinate analysis (PCoA), and significance was tested using the Wilcoxon rank-sum test and by analysis of similarity (ANOSIM). Functional differential abundance was assessed by a linear discriminant analysis effect size (LEfSe) algorithm (Segata et al. 2011). False discovery rate (FDR) p-value correction was applied to all tests in this study.

## RESULTS

### Bacterial community profiling

#### Comparison of seed epiphyte microbiomes of fresh and stored seed

A total of 2443 OTUs were identified in the dataset following sequence quality checking and filtering. All OTUs which were assigned a taxonomy were bacterial, with no archaea identified. Of these, the majority of OTUs were found to be unique to fresh seeds (42.5%), and few OTUs were identified to be unique to a specific location (3.1%). A set of core OTUs were observed that were present across all fresh samples, these core taxa while only representing ∼20% of recovered OTUs were important contributors in terms of overall relative abundance, with 71.8% of quality filtered reads belonging to these bacteria (Figure 2).

**Figure 2:**
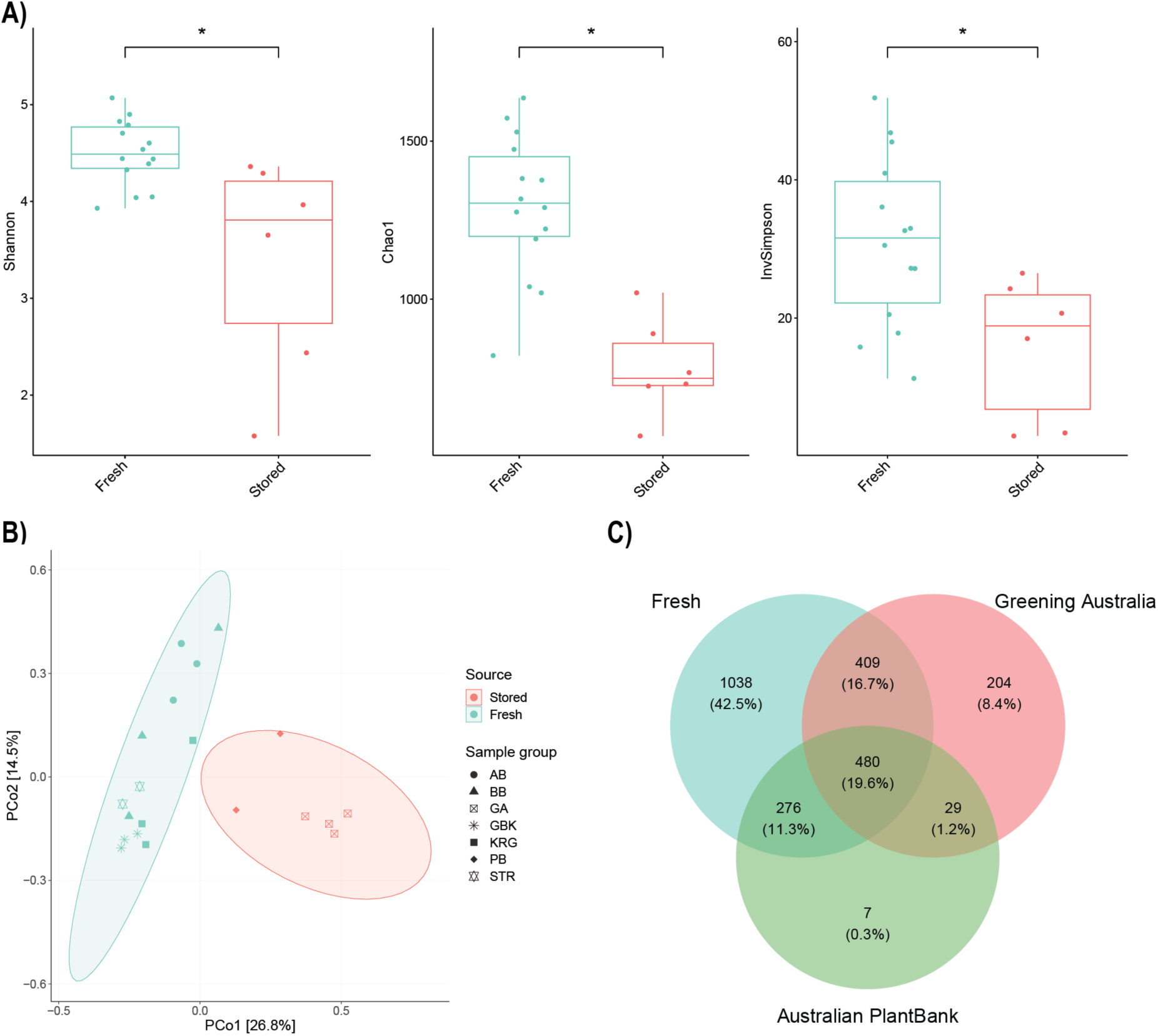
*Acacia ulicifolia* seed-associated microbial community diversity and composition represented by **(A)** alpha **(B)** beta diversity between stored and fresh seed accessions. Diversity of microbial communities assessed based on Shannon diversity indices, Chao1 species richness, and Inverse Simpson evenness according to storage status **(A)**. Microbiome community composition was assessed based on storage status (colour) and sampling location (points represented by different symbols) using a Bray-Curtis dissimilarity matrix **(B)**. Venn diagram **(C)** indicating the number and proportion of unique and shared OTUs between the seed microbiomes of samples from each seed source.

The epiphytic bacterial communities on *A. ulicifolia* seeds from freshly collected and stored seed samples were compared, looking at both bacterial diversity and community composition. Stored samples were found to be significantly lower in bacterial diversity (*p* < 0.001), species richness and evenness (*p* < 0.001) than fresh samples (Figure 2A). Provenance factors influencing seed microbiome composition were examined using Principal Coordinate Analysis (PCoA), which also showed that stored and fresh seed bacterial communities were distinct, clustering separately (Figure 2B). Differences in overall composition were confirmed to be significant according to a Wilcoxon rank-sum test (*p* < 0.001) (). ANOSIM results confirmed storage condition to be a principal driver of variation (*R* = 0.8383, *p* < 0.001). Sampling location was also found to contribute to variance between fresh samples to a significant yet lesser extent (*R* = 0.2939. *p* = 0.0170). In contrast, within freshly collected samples, alpha diversity (Shannon) and species richness (Chao1) were comparable across all locations surveyed (*p* > 0.05) and no significant difference in evenness was identified (InvSimpson, *p* = 0.98)

#### Community composition of A. ulicifolia seed microbiomes

Of the 2443 OTUs identified from the seed epiphytic communities, there appeared to be a core set of OTUs that were consistently found in all freshly collected seed samples across the different sampled locations. These OTUs were affiliated with the phyla *Alphaproteobacteria*, *Gammaproteobacteria*, *Acidobacteriota*, and *Actinobacteriota* and in every fresh sample these groups together accounted for approximately 50% of the total relative abundance. This core set was comprised of OTUs that are currently not well characterised below genus level, including an undescribed *Rhizobiales* genus within the *Beijerinckiaceae* family (1174-901-12), and representatives of *Sphingomonas*, *Methylobacterium-Methylorubrum*, *Allorhizobium-Neorhizobium-Pararhizobium-Rhizobium* (hereafter *Rhizobium*), and *Terriglobus.* The remainder of OTUs in fresh seed microbiota consisted of a diverse assortment of genera each of which had low individual abundance (< 5%). Interestingly, a sizeable proportion of these included OTUs assigned to genera for which there are currently no cultured representatives in the Silva v. 138 database (Figure 3C). In contrast, microbiomes of stored seeds contained far fewer low-abundance taxa. Instead, the microbiome of GA seeds largely consisted of *Bacillus* and *Paenibacillus,* whereas PB seeds were predominantly comprised of *Erwinia, Pantoea*, and *Pseudomonas*. Additionally, *Massilia* and *Curtobacterium* were abundant genera common to both sources of stored seed.

**Figure 3:**
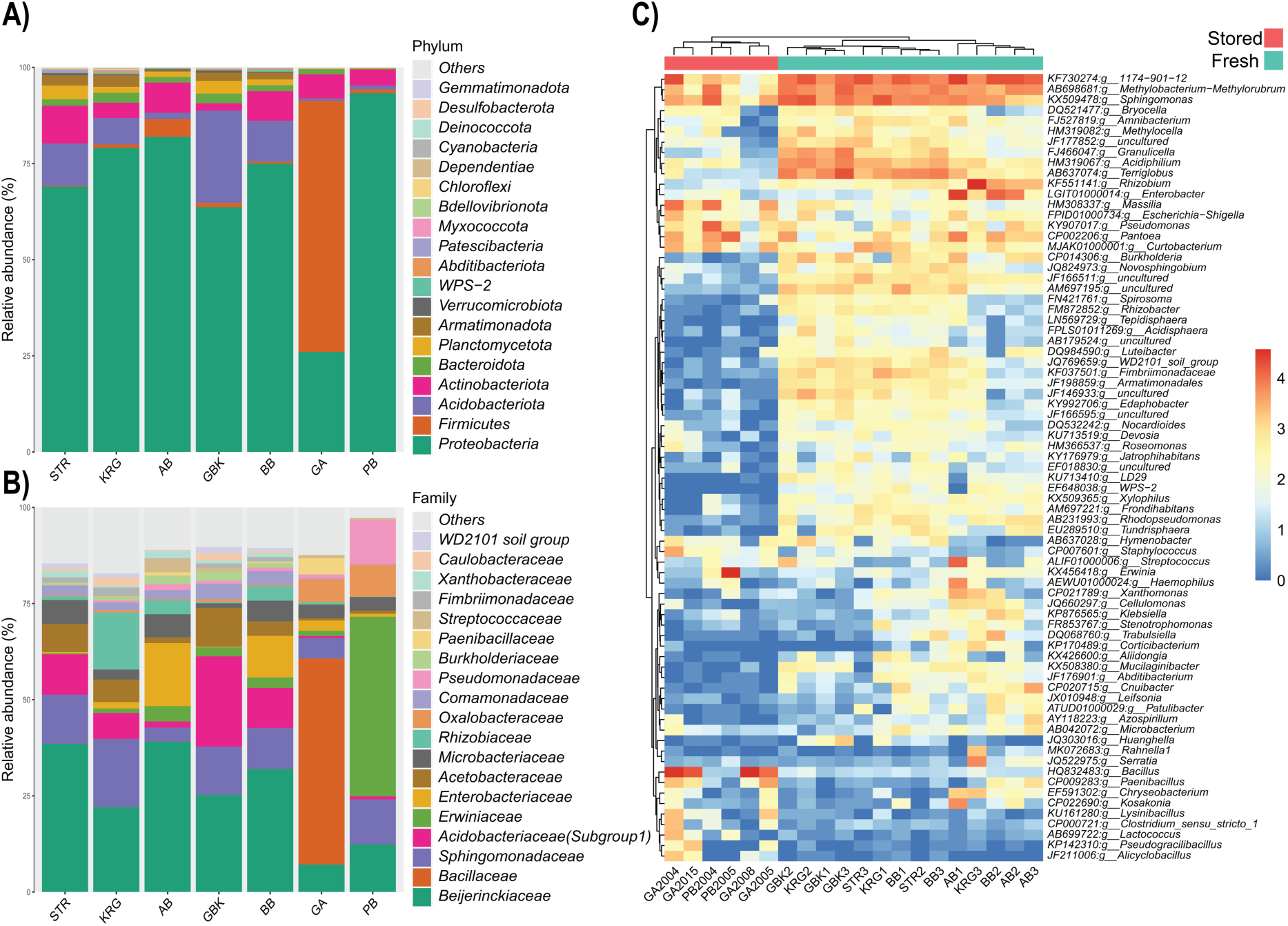
Relative composition of the *A. ulicifolia* bacterial community at **(A)** phylum and **(B)** family taxonomic level for each sample location. **(C)** Bacterial abundance heatmap profile of the top 75 most abundant genera in *A. ulicifolia* seed-associated bacterial communities between stored (red) and fresh (blue) samples.

### Functional profiling of culturable bacteria from fresh and stored *A. ulicifolia* seed

A total of 243 morphologically distinct bacteria were isolated from *A. ulicifolia* seed surfaces (224 from fresh samples and 19 from stored samples). Screening for three plant growth-promoting (PGP) phenotypes (nitrogen fixing, phosphate mobilisation, or IAA production) indicated most isolates possessed at least of one these traits (72.8% of fresh samples 36.7% of stored samples). The main bacterial morphotypes from this collection (70 from fresh seed accessions and 11 from stored seed) were subject to further analysis to determine taxonomy and extend the PGP trait profiling. The taxonomic profiling, performed via 16S rRNA gene sequencing, showed that this isolate collection included representatives of 28 different genera, with *Curtobacterium*, *Pseudomonas, Microbacterium*, *Luteibacter*, *Xanthomonas*, and *Methylobacterium* among the most abundant groups observed (Figure 4). Of these 81 bacterial isolates, 39.5% possessed ACC deaminase activity, 54.3% produced siderophores, 46.9% produced IAA, 14.8% solubilised inorganic phosphate, while 60.4% displayed biological nitrogen fixation capacity. Most isolates (87.7%) possessed at least two traits (Figure 4). Notably, one isolate recovered from fresh seed, a representative of the genus *Leclercia,* possessed all traits screened.

**Figure 4:**
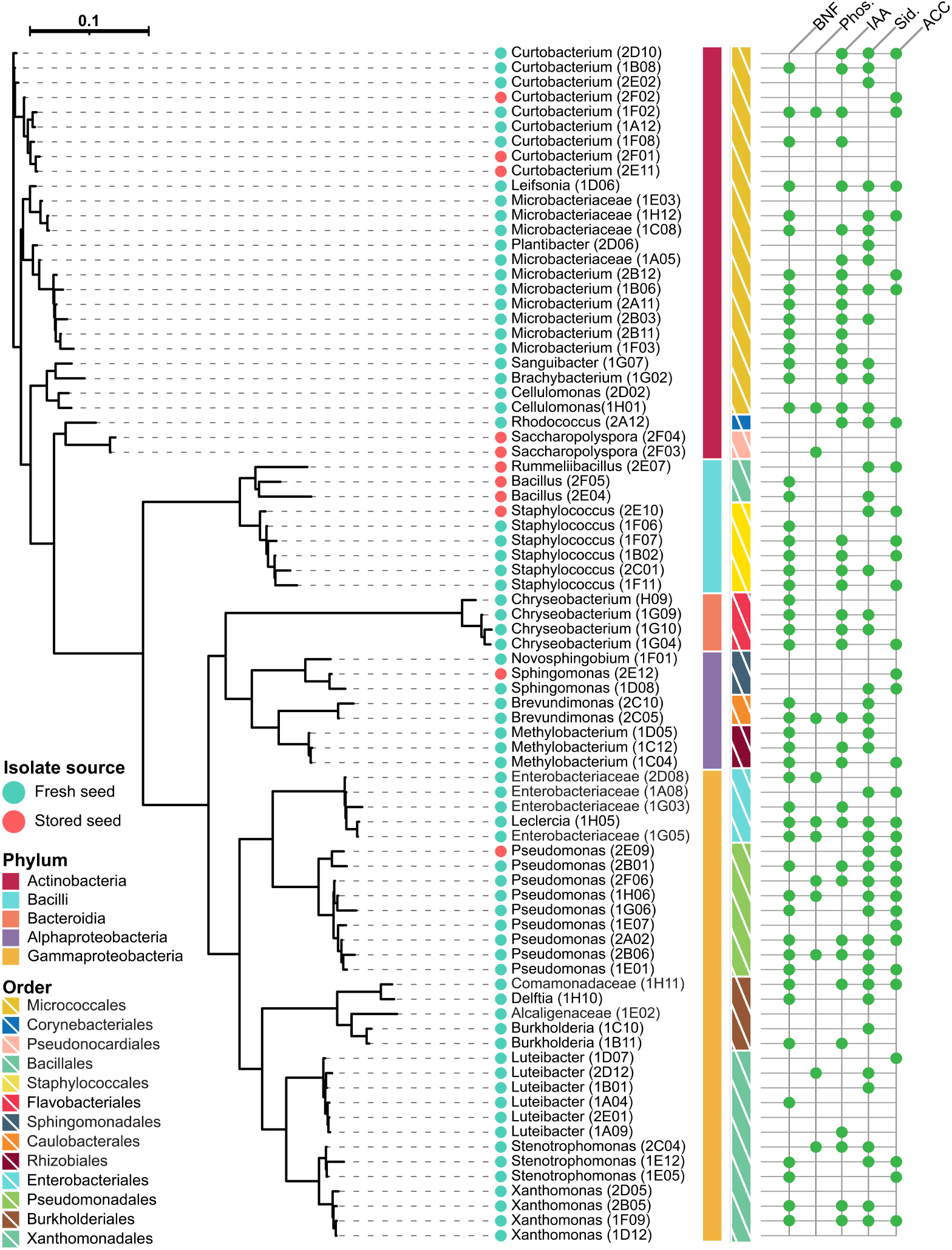
Phylogenetic analysis inferred from maximum likelihood (RAxML) analysis of 16S rRNA gene sequences from bacterial isolates recovered from *A. ulicifolia* seeds. Taxonomic assignment based on classification in the SILVA (v. 138) database using the Alignment, Classification and Tree (ACT) tool. Putative plant growth promoting functions for each isolate determined *in vitro* indicated by green circles for each trait tested (BNF = biological nitrogen fixation, Phos. = phosphate solubilisation, IAA = Indole-3-acetic acid production, Sid. = Siderophore production, ACC = 1-aminocyclopropane carboxylic acid deaminase activity). Isolates recovered from stored and fresh seeds are indicated by red and blue circles respectively.

### Predicted functional capacity of the bacterial seed microbiome

Microbiome functional predictions were performed using 16S rRNA gene community sequencing data and Tax4Fun2 to generate trait profiles. Predicted functional profiles of stored and fresh seed were compared to identify potential differences, focusing on traits of relevance to microbe-plant interactions, using the PLaBAse PGPTO database to identify pathways relevant to plant hosts. A set of 40 functions were found to be significantly enriched/depleted between stored and fresh seeds (*p* < 0.05) based on LEfSe differential abundance analysis. Fresh seed microbiota were found to be enriched for functions associated with nitrogen fixation, phosphate enrichment, bacterial antagonism, plant resource utilisation, and resistance to mercury and arsenic, compared to stored seeds. In contrast, the functional profile of stored seeds was enriched with traits associated with fungal antagonism, potassium enrichment, thiamin biosynthesis, and the *ter* operon (a set of genes associated with tellurite resistance) (Figure 6). No significant difference in trait profiles was identified between different fresh sampling locations, or different seed storage facilities (*p* > 0.05) following *p*-value adjustment.

### Estimates of seed-associated bacterial cell densities

To determine if seed surfaces from different samples hosted bacterial communities of different density, flow cytometric counts of DNA-stained cells 1.0 µm – 6.0 µm in size (the size range of most culturable bacterial cells) were calculated per gram of seed and log transformed (Figure 5A). For all freshly collected samples, as well as GA-stored seed, a sizable cell population was observed in the seed wash (> 10^7^ cells per gram washed seed). However, stored seed from PB samples had significantly (*p* < 0.001) lower cell counts than all other sample groups, with considerable variation between counts for specific PB samples. Flow cytometric estimates of bacterial populations within each seed wash were also observed to be fairly consistent for samples from most locations, with the exception of PB samples.

**Figure 5:**
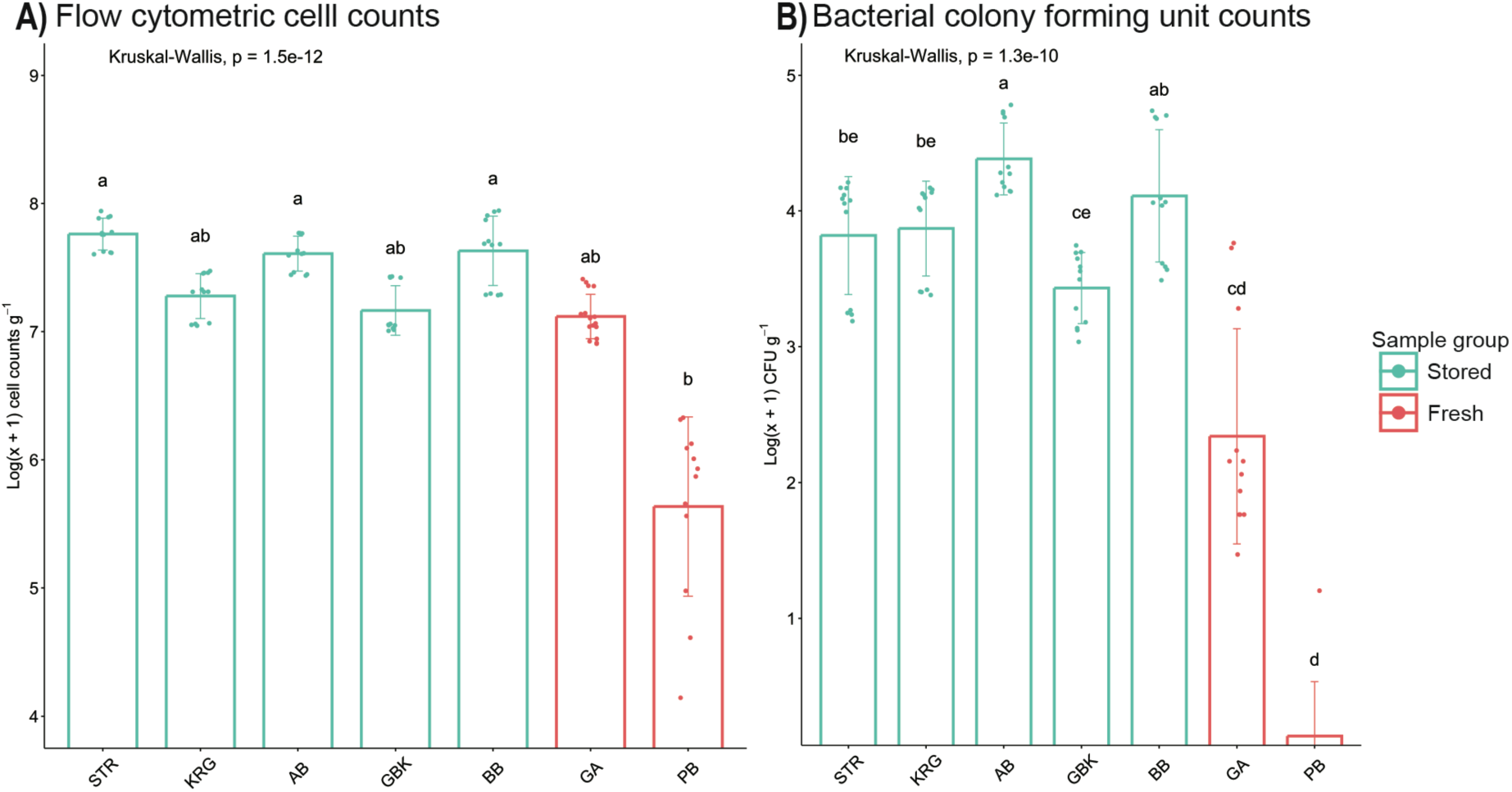
Microbial cell counts of *A. ulicifolia* seed epiphytes determined by **(A)** flow cytometry and **(B)** colony forming unit counts. Counts are reported as mean (n = 12) values for each fresh seed sampling location (blue/green) or stored sample group (red). Significant differences, as determined by Kruskal-Wallis tests, are denoted unshared letters.

**Figure 6:**
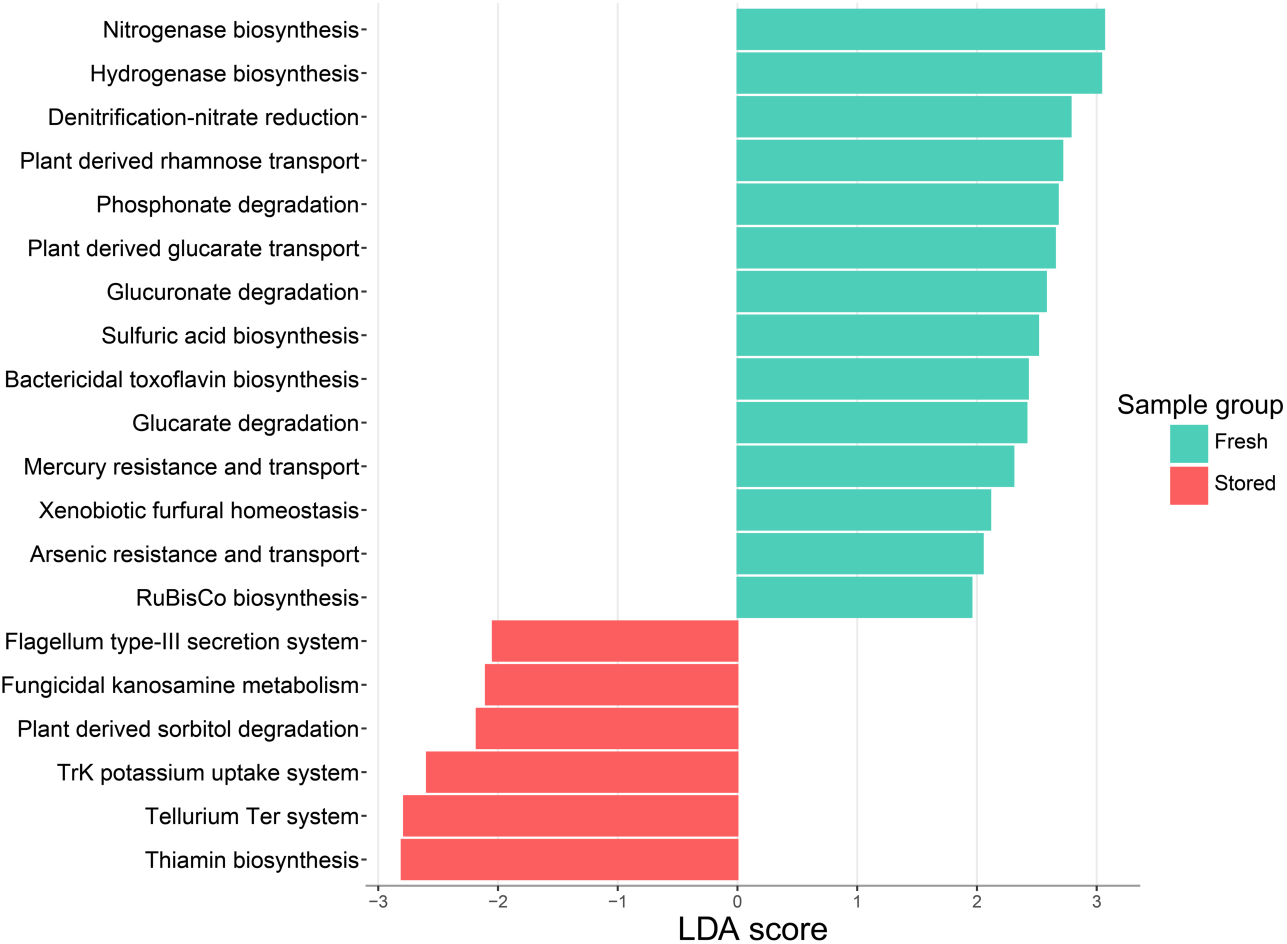
Differentially-abundant plant growth-promoting functions predicted by taxonomic inference using Tax4fun2, as determined by linear discriminant analysis (LDA). Functions classified by PLaBAse plant growth promoting trait ontology database.

The number of culturable bacterial colonies recovered from each seed surface (expressed as CFU per gram of seed) was also counted following 10 days of incubation, as another measure of bacterial density (Figure 5B). Bacterial colonies were recovered from all samples except for the stored sample PB1988 where no growth was observed following the incubation period. When compared to flow cytometric counts, CFU estimates displayed considerable variation within and between sample groups. Counts of CFU were also consistently several-fold lower than flow cytometric cell counts, likely reflecting the high proportion of environmental bacteria that are unculturable under standard laboratory conditions.

Culture analysis reflected flow cytometry results in that the number of bacterial colonies recovered from seeds tended to be lower for stored than for freshly collected samples, with PB samples in particular having very few bacterial colonies recovered. Pairwise analysis was performed and showed there was a significant difference in average CFU per gram of seed between some sample groups (*p* < 0.001, Fig 5B), with PB samples significantly lower than any of the fresh samples.

## DISCUSSION

High quality seed from native species is an essential resource for restoration and conservation of plant diversity the world over. Seed-associated bacteria can play a demonstrated role in ensuring seed quality in relation to germinability and viability (Berg and Raaijmakers 2018, Rodriguez et al. 2020, Mertin et al. 2023) but seed storage under standard seed bank conditions may affect the load and composition of seed microbiomes, impacting quality. We show that the epiphytic microbiome of seedbank-stored native seeds hosted less diverse communities of bacteria, coinciding with a significantly reduced total bacterial population relative to fresh, field-collected seeds. Bacterial community composition also differed substantially between stored and field-collected seeds, with many taxa abundant in fresh seeds, including members with plant-beneficial functional traits, absent in stored seeds. Our results have implications for the successful downstream use of seeds in nature repair programs, particularly where altered or reduced bacterial composition affects germination and early-stage growth.

### Understanding native seed microbiomes

While much of the research on seed microbiomes has focussed on agricultural species, those of native seeds are beginning to receive research attention. Cross-taxon studies have identified bacterial groups common across seed microbiomes and have shown that native seeds tend to host more diverse microbiomes compared to seeds from agricultural plant species (Wasserman et al. 2019, Chandel et al. 2022). Many of the genera observed to be abundant in seed microbiome of our study species are common to a diverse suite of plant species. For instance, reviews of seed microbiome research emphasise *Pantoea*, *Sphingomonas*, *Methylobacterium*, *Bacillus*, *Curtobacterium* and *Pseudomonas* as prominent seed endophytes and epiphytes in both native and domesticated plant lineages (Simonin et al. 2022, Truyens et al. 2015, Wasserman et al. 2019).

We show that *Sphingomonas*, *Methylobacterium*, *Rhizobium*, *Curtobacterium,* and *1174-901-12* (a member of *Rhizobiales* belonging to an undescribed genus) were consistently key contributors in terms of relative abundance in fresh seed, with *Pseudomonas* also important in seeds from specific sites based on 16S rRNA sequencing. Culture-based work also regularly recovered isolates of *Microbacterium*, *Luteibacter*, *Staphylococcus* and *Cellulomonas*, but did not recover some core microbiome components, such as 1*174-901-12*. Interestingly *A. ulicifolia* seeds hosted significant populations of subdivision 1 *Acidobacteria* including *Terriglobus*, *Granulicella*, and *Bryocella*. Their presence on seeds is unusual, as these genera, in addition to *Acidiphilium*, typically reside in mineral or soil environments and are only reported to colonise aboveground plant surfaces in a small number of studies (Noble et al. 2020, Hakobyan et al. 2023). Previous research has noted the abundance of these taxa in phyllosphere microbiomes correlates with seasonal changes in precipitation (Li et al. 2022) with suggestions they may be deposited on aerial plant surfaces by rain without necessarily colonising the host (Mechan Llontop et al. 2021). Whilst other prominent members of the *A. ulicifolia* seed microbiome including *1174-901-12*, *Sphingomonas*, *Pseudomonas*, and *Methylobacterium* have also been reported to be transmitted by rain, these genera are prominent colonisers of mature aerial and internal compartments in diverse plant species (Espenshade et al. 2019, Anguita-Maeso et al. 2020, Ares et al. 2021, Dreyling et al. 2022).

Overall, our results suggest that some fraction of the *A. ulicifolia* seed microbiome is likely dynamic, with fresh seed microbiomes comprised of transient or locally adapted bacterial taxa in addition to specifically adapted *A. ulicifolia* core seed microbiota. Undoubtedly, repeated analysis of samples of seed at the same locations through time would contribute to a clearer picture of seed epiphytic microbiome dynamics. Such work is important given knowledge of epiphytic microbial communities on seed remains limited. Their composition may shift considerably through time and is determined by the local environment, contrasting with endophytes which are thought to establish long-term symbioses with hosts (Abdelfattah et al. 2023). Significant variation in epiphyte communities along spatial and temporal gradients has been identified in seed of agricultural *Brassica* and *Triticum* cultivars, indicating that the microbiome is determined almost exclusively by environmental factors (Barret et al. 2015, Klaedtke et al. 2016). However, other studies report consistent populations of shared microbiota within the same hosts between distant populations (Links et al. 2014, Morales-Moreira et al. 2021).

### Differences in stored and freshly collected seed microbiomes

Although seed processing, drying, and low-temperature storage remain necessary to preserve seed viability, our results highlight that important microbial symbionts may be lost in stored seed. We identified substantial reductions in bacterial load and community diversity associated with stored seeds, driven primarily by the absence of rare and low-abundance taxa compared to freshly collected counterparts. This is similar to what was previously noted in work looking at artificially stored soybean endophytes (Chandel et al. 2021). Although studies examining the effects of postharvest seed storage remain scarce, available evidence suggests that bacteria with specific adaptive strategies to desiccation and temperature stress are selected for during the storage process. The present study and previous literature examining stored seed microbiota indicate stored seed microbiomes may be enriched in *Bacilli* and *Gammaproteobacteria* groups possessing traits such as sporulation, peptidoglycan cell walls, or secretion of extracellular polysaccharides, known to improve tolerance to temperature and desiccation stresses (Beringer et al. 2018). This most commonly includes strains of *Pantoea*, *Erwinia*, and *Bacillus*, reported as abundant members of both endophytic and epiphytic stored seed communities (Chandel et al. 2021, Dutta et al. 2022), likely a consequence of reduced moisture content also occurring within stored seed tissues.

Certain members of the core epiphytic community of *A. ulicifolia, 1174-901-12*, *Sphingomonas*, and *Methylobacterium* common in fresh seed were found to persist in stored seed samples, albeit in reduced relative abundance. These genera have been reported to together form pioneer biofilms on a range of surfaces susceptible to desiccation stress such as photovoltaic panels and roof tiles (Moura et al. 2021, Romani et al. 2019), which possibly also contributes some degree of protection during seed storage. However, it should be acknowledged that, as few isolates of *Sphingomonas* or *Methylobacterium* were recovered from stored seed, it is possible 16S rRNA gene-based community profiling reflects dormant unculturable bacteria or nonviable cells.

Differences in bacterial load and composition were also observed between each of the two storage facilities from which samples were sourced, suggesting that differences in handling practices may also shape stored seed microbiomes. It is worth noting that there were differences in temperature preferences of the dominant taxa from different locations (psychrotolerant in PlantBank and mesophilic in Greening Australia) consistent with 4°C storage vs room temperature storage respectively. Past work has shown temperature can influence the composition of stored seed microbiota (Chandel et al. 2021). However, it is likely that multiple environmental parameters, including temperature, desiccation, and variations in seed handling together with storage time all contribute to observed differences between organizations. To dissect the contributions of each of these factors to stored seed microbiomes, manipulative experiments examining each independently are required. While we cannot comment on the microbiome of seeds before storage in this study, the work presented here supports previous assertions that storage may result in substantial reductions in diversity in the seed microbiome.

### Ecological significance of changes in seed microbiota

In this study, we isolated numerous bacteria from freshly collected *A. ulicifolia* seed surfaces possessing well-established plant growth-promoting (PGP) phenotypes. This included evidence of ACC deaminase activity and IAA production, traits that have been shown to improve tolerance to drought and salinity stress across a wide range of plant species (Hone et al. 2021, Ratnaningsih et al. 2023). Many of our isolated strains also show capacity for N-fixation, siderophore production and phosphate solubilisation, mechanisms which have been demonstrated to overcome nutrient deficiency (Borken et al. 2016, Shao et al. 2021), pathogenic invasion (Matsumoto et al. 2021), and more generally enhance host productivity (Abdelfattah et al. 2021, Rodriguez et al. 2020, Verma et al. 2018). Bacterial isolates possessing these traits, such as those identified in this study, may also synergistically interact leading to emergent properties with improved plant growth-promoting capacity (Yang et al. 2021), although we did not examine this. While stored seeds were not entirely lacking in isolates with PGP traits, only a small number of such isolates were recovered from stored seed, and few of these showed multiple beneficial traits. Stored seeds were also found to host bacterial communities possessing fewer traits relating to the utilisation of plant resources compared to fresh seed, functions known to influence colonisation success during seedling emergence (Torres-Cortés et al. 2018). As microbes on seed surfaces are important in shaping seedling microbiomes (Arnault et al. 2024, Moroenyane et al. 2021), the reduced diversity of PGP bacteria associated with stored seeds may reflect microbiome changes which have the potential to influence the health and survivorship of seedlings produced from stored seed. However, the true impact of storage-induced changes to the microbiome cannot be adequately assessed without further studies *in planta*.

As degraded ecosystems of interest to restoration practitioners are typically characterised by soils depleted in nutrients and microbial diversity (Hart et al. 2020), seed microbiota may represent a particularly important source of essential symbionts available to the developing seedling (Walsh et al., 2021, Moroenyane et al. 2021). Research indicates that changes in low-abundance taxa, such as those reported in this study, may lead to substantial changes in plant growth-promoting activity in microbial communities (Shao et al. 2021, Tom et al. 2023, Pan et al. 2024). Furthermore, although our understanding of conserved symbionts in *A. ulicifolia* is limited, the presence of core genera which showed high relative abundance across all or most samples implies these taxa may play an important ecological role in host functioning. Most significantly, *Rhizobia* and related members of *Beijerinckiaceae* comprise taxa shown to stimulate nodulation as well as engage in associative symbiotic N-fixation with host plants (Tamas et al. 2014, Borken et al. 2016). In many species of *Acacia* they are known to be essential to healthy functioning (Barrett et al. 2015, Thrall et al. 2005), and so their absence in stored seeds is concerning. Overall, it is reasonable to suggest that the seed banking process may reduce seed associated bacterial diversity, truncating the number of potentially important symbionts available to developing seedlings. In light of this, our work supports growing calls for further consideration of the seed-associated microbiome during the storage process (Mertin et al. 2023, Berg and Raaijmakers 2018).

Synthesising data from 16S rRNA gene-based community characterisation, flow cytometric cell counts, culture-based screening and phenotypic assays we have characterised the structure, function and load of the epiphytic seed microbiome of *A. ulicifolia* sourced fresh from native populations and from seed banking organizations. We demonstrate the epiphytic microbiome of freshly collected *A. ulicifolia* seeds consist of a diverse range of bacterial taxa comprising both core and rare taxa, including many strains with plant growth-promoting capacity. While the full significance of the seed microbiome to native species’ health and success in restoration work requires further investigation, results presented in this study suggest that some of these potentially beneficial microbes may be lost during storage. We demonstrate that the surfaces of *A. ulicifolia* seeds obtained from seedbanks support fewer and less diverse bacterial populations when compared to natural seed populations. Changes to the microbiome composition of stored seeds observed in this work reflect what has been reported in the small number of manipulative experiments published previously on seed storage and microbial composition for other plant species. Conventional seed processing and storage practices most likely select for members of the bacterial community with adaptation strategies that facilitate survival under low temperature and humidity conditions. Further research is required to determine the impact of these changes of the microbiome *in planta*, and the flow-on effects to restoration success.

#### DATA AVAILABILITY

The sequencing data from this study have been deposited under NCBI BioProject ID PRJNA1134362.

#### AUTHOR CONTRIBUTIONS

Conceptualisation of study: DHR, RVG, MMM, SGT

Experimental investigation and validation: DHR, VR

Data analysis: DHR, VR

Project management and supervision: RVG, SGT

Writing the original draft of the manuscript: DHR

Reviewing and editing the manuscript: DHR, MA, RVG, MMM, SGT

## ACKNOWLEDGEMENTS

This work was funded by the Australian Research Council Linkage Project grant LP200200688 to RVG, SGT and MMM. The authors would like to acknowledge the contribution of Paden Willson from Greening Australia, and Katherine Thompson from the Australian PlantBank for supplying seeds used in this study.

